# Spatial Transcriptomics of TNBC tumours and corresponding lymph node metastasis reveals immune hubs driven by TNF/NF-κB signalling of TMSB4X and CD74 expressing cells

**DOI:** 10.64898/2026.03.15.711865

**Authors:** Adrian Kacperczyk-Perdyan, Urszula Ławrynowicz, Maciej Jaśkiewicz, Anna Kostecka, Katarzyna Chojnowska, Mikołaj Koszyński, Marcin Jąkalski, Michał Bieńkowski, Natalia Filipowicz, Agnieszka Misztak, Tomasz Nowikiewicz, Łukasz Szylberg, Marta Drzewiecka, Arkadiusz Piotrowski, Jakub Mieczkowski

## Abstract

**Purpose:** Lymph node metastasis is a critical prognostic factor in triple-negative breast cancer (TNBC), but the spatial organization of signalling networks driving metastatic colonization and persistence remains poorly defined. Understanding these networks may reveal therapeutic vulnerabilities in metastatic TNBC.

**Methods:** We applied Visium spatial transcriptomics to paired primary tumours and lymph node metastases from four TNBC patients. Spatial gene expression profiles were analysed using trajectory inference, cell-type deconvolution, and cell-cell communication mapping. Results were validated in independent single-cell RNA sequencing datasets from TNBC tumours and lymph node metastases.

**Results:** TMSB4X and CD74 emerged as key drivers of metastasis-associated transcriptional programs, defining spatially distinct immune communication hubs enriched in myeloid, stromal, and endothelial cells. These hubs preferentially activated NF-κB/TNF signalling alongside PI3K-Akt and Rap1 pathways. In primary tumours, NF-κB/TNF signalling was confined to localized immune hubs, whereas lymph node metastases exhibited widespread signalling across cellular compartments with additional IL-1 pathway activation. This spatial rewiring coincided with endothelial cells assuming central coordinating roles in metastatic lesions. Independent validation confirmed myeloid-endothelial crosstalk as a conserved feature of TNBC, with TMSB4X-CD74 programs enriched in clinically relevant immune and vascular subpopulations.

**Conclusion:** TNBC lymph node metastases display spatially expanded inflammatory networks centred on TMSB4X-CD74 immune hubs. Metastatic progression involves coordinated myeloid-endothelial signalling, suggesting these pathways as potential biomarkers and therapeutic targets. Disrupting inflammatory and vascular communication networks may offer a strategy to prevent metastatic persistence and overcome therapy resistance in TNBC.

## INTRODUCTION

Breast cancer (BC) is the most frequently diagnosed cancer in women worldwide and a leading cause of cancer-related death, largely due to metastasis [1]. Axillary lymph nodes (LNs) represent the first site of tumour dissemination and LN involvement is associated with poor prognosis and recurrence, yet the molecular and spatial organization of LN metastases remains insufficiently characterized [2]. This knowledge gap hampers the development of effective therapies targeting metastatic disease.

BC is a heterogeneous disease comprising distinct molecular subtypes, including luminal A, luminal B, HER2-enriched, and triple-negative breast cancer (TNBC) [3]. TNBC accounts for 15–20% of BC cases and is defined by the lack of ER, PR, and HER2 expression, aggressive clinical behaviour, early relapse, and limited therapeutic options [4]. Pronounced intratumoral heterogeneity further contributes to treatment resistance, underscoring the need to better understand the molecular drivers and microenvironmental interactions underlying TNBC metastasis [5].

Spatial transcriptomics (ST) enables the analysis of gene expression within intact tissue architecture, preserving spatial context and cell–cell interactions that are lost in bulk or single-cell approaches [6, 7]. The Visium Spatial Gene Expression (VSGE) platform allows transcriptome-wide profiling of FFPE samples, providing access to clinically annotated archival material. Despite the importance of LNs in early metastatic spread, it remains unclear whether the signalling and cellular networks present in primary TNBC tumours are preserved or rewired in LN metastases.

Here, we applied ST to matched primary tumours and LN metastases from four TNBC patients. By integrating spatial gene expression data with pathological assessment and publicly available single-cell RNA sequencing datasets, we sought to elucidate spatially resolved regulatory networks driving metastatic progression and to identify potential therapeutic vulnerabilities in TNBC LN metastases.

## MATERIALS AND METHODS

### Tissue samples

Tissue specimens were provided by Oncology Centre-Prof. Franciszek Łukaszczyk Memorial Hospital in Bydgoszcz. All experimental procedures were conducted in accordance with Independent Bioethics Committee for Scientific Research at Medical University of Gdansk (consents No. NKBBN/563/2018 and NKBBN/564-108/2022). Paired primary tumours (PT) and matched lymph node (LN) metastases from four TNBC patients were included in the study.

### FFPE RNA Quality Check

RNA quality was evaluated by bulk RNA extraction from FFPE tissue sections using the RNeasy FFPE Kit (Qiagen). The DV200 metric, defined as the proportion of RNA fragments longer than 200 nucleotides, was used to assess RNA integrity. Only blocks with DV200 ≥ 60% were selected for further analyses, in line with recommended thresholds for robust Visium FFPE assay performance (Supplementary Table 1).

### Cryosection

FFPE blocks from PT and LN metastases were sectioned at the Department of Pathomorphology, Medical University of Gdansk. Tissue sections were mounted onto the Visium (VSGE) slides according to the manufacturer’s guidance. Slides were incubated for 3 hours at 42 °C and dried overnight at room temperature.

### H&E staining, and brightfield imaging for ST

Deparaffinization and H&E staining were performed following the Visium Spatial for FFPE Demonstrated Protocol (CG000409 Rev. D). Brightfield imaging was carried out to confirm tissue integrity and proper placement on capture areas. After imaging, slides were prepared for Probe hybridization, performed overnight at 50 °C in accordance with the VSGE for FFPE User Guide (CG000407 Rev. E).

### VSGE library construction and sequencing

Probe ligation, release, and library construction followed the Visium FFPE protocol (CG000407 Rev. E). PCR cycle number was determined by qPCR (LightCycler® 480; Cq range: 9.2–9.8), with final cycles set as Cq + 2. Libraries were indexed using 10× Genomics Dual Index TS Set A and quantified by qPCR (KAPA Library Quantification Kit). Sequencing depth was calculated as: (tissue coverage fraction) × 5,000 spots × 25,000 read pairs per spot. Pooled libraries were sequenced on Illumina NextSeq 550 (Read 1: 28 cycles; i7/i5 indices: 10 cycles each; Read 2: 50 cycles).

### VSGE data processing

Raw sequencing data (BCL files) from four PT-LN pairs were converted to FASTQ format using Space Ranger (v2.0.0). Reads were aligned to the human reference genome (GRCh38) using STAR, and spatial feature counts were quantified with Visium Human Transcriptome Probe Set v1.0. Loupe Browser 8.0 was used for alignment visualization and quality control.

### Data preprocessing, clustering, and differential expression

Spatial transcriptomic data were processed in Seurat (v5.3) using the *Load10X_Spatial* function. Low-quality spots were excluded based on feature counts (<100 or >12,000) or mitochondrial gene content (>15%). Data were normalized using *NormalizeData* and *SCTransform* functions with default parameters. Highly variable features were identified using variance-stabilizing transformation (VST), and data were scaled with regression of mitochondrial gene content. Dimensionality reduction was performed using PCA followed by UMAP. Nearest neighbours were computed using the first 20 principal components based on ElbowPlot, and clustering was performed using a shared nearest-neighbour (SNN) algorithm at a resolution of 0.8. Differentially expressed genes (DEGs) were identified using *FindAllMarkers* (only.pos = TRUE, test.use = “MAST”, min.pct = 0.25, logfc.threshold = 0.25), retaining genes with P < 0.05 [8]. MAST is a flexible framework for single-cell RNA-seq that models both the probability of expression and the expression level, enabling supervised differential expression analyses and unsupervised exploration of co-expression patterns [8].

### Cell type deconvolution and CNV analysis

Robust Cell Type Decomposition (RCTD) was performed using a single-cell RNA reference deposited in GEO under accession code GSE180286, comprising two TNBC patients with matched LN metastases [9]. Single-cell data were processed in Seurat using filtering thresholds based on gene count quantiles (2.5–97.5%) and mitochondrial content (<5%). Single-cell data were filtered based on gene count quantiles (2.5–97.5%) and mitochondrial content (<5%), clustered at resolution 2, and annotated using canonical marker genes. Copy-number variation analysis was conducted using *inferCNV* (v1.24.0), with B cells as the normal reference to distinguish malignant epithelial cells [10]. RCTD was run in doublet mode, and deconvoluted cell-type proportions were visualized using *SpatialFeaturePlot*.

### Trajectory, enrichment, and communication analyses

Trajectory analysis was conducted using Monocle3 [11], with pseudotime ordering and identification of trajectory-associated genes using *graph_test* (principal_graph neighbourhood, q < 0.05). Shared trajectory drivers were defined based on the top 0.5–2% of Moran’s I statistics. Functional enrichment analysis of DEGs was conducted using KEGG and visualized with clusterProfiler [12], with Benjamini-Hochberg (BH) adjusted P < 0.05 considered significant. Cell–cell communication was analysed using CellChat (v2.2.0) to infer ligand– receptor interactions and pathway-level signalling [13]. Validation was performed using independent TNBC scRNA datasets (GSE180286 [9] and GSE176078 [14]) by assessing TMSB4X–CD74 expression correlations.

### Data availability

This study used publicly available data repository (Gene Expression Omnibus https://www.ncbi.nlm.nih.gov/geo/. Detailed description of each dataset used can be found in methods section of the manuscript. The R code used in the analysis was published on GitHub repository (https://github.com/jakubmie/).

## RESULTS

### Spatial mapping of cell composition in TNBC tumours and metastases

We applied VSGE profiling to four paired samples of TNBC PT and matched LN metastases (Figure 1A). All tissue specimens (n=8) were examined and delineated by an experienced pathologist (Figure 1B; Supplement Figure 1A-C). Patient clinical characteristics are summarized in Supplement Table 1.

**Figure 1.**
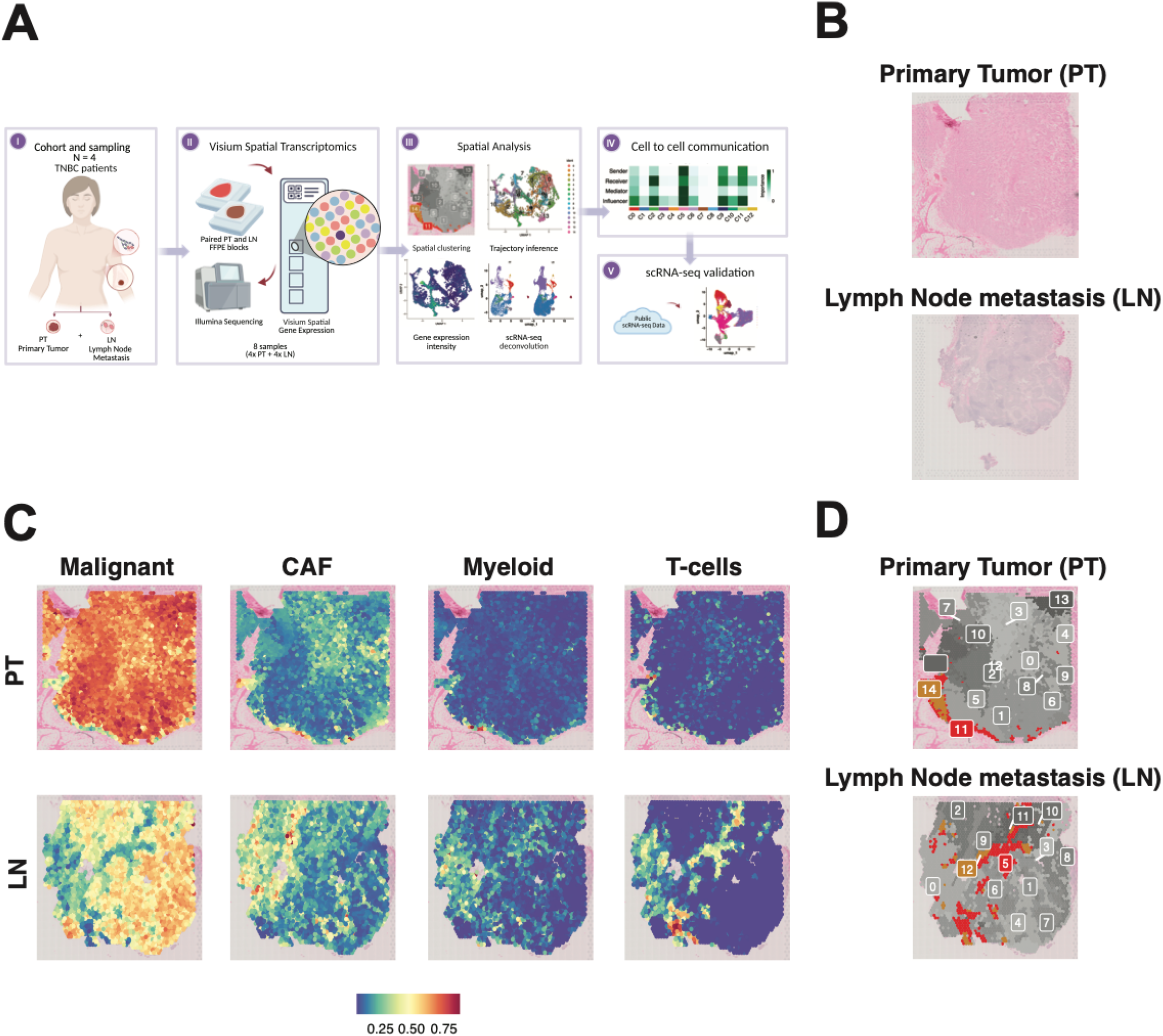
Spatially resolved primary tumour and lymph node metastasis profiles identify transcriptionally distinct clusters. Data present for patient 1 (FG1XW) out of four examined patients. Graphs for other patients are present in the Supplement. A) Schematic presentation of the study design. B) FFPE tissue slides used for Visium Spatial Gene Expression experiment. C) Spatial distribution of cell-type probability of major cell types in each spot of Visium slide. Probabilities were assigned based on RCTD for which we used previously created RNA reference. Based on RCTD probabilities two major cell-types present are malignant cells and cancer-associated fibroblasts. D) Seurat clustering used for downstream analysis. Marked clusters (orange, red) are clusters with the highest expression of CD74 and TMSBX4 genes (for details continue to Figure 2). Figure legend: CAF – cancer associated fibroblast; FFPE - Formalin-Fixed Paraffin-Embedded; LN – lymph node; N – Number; PT – primary tumour; RCTD - Robust Cell Type Decomposition; scRNA-seq – single-cell RNA; TNBC – Triple Negative Breast Cancer.

To examine the cell composition in each spot, we performed the RCTD deconvolution, Seurat clustering and DE analysis (see methods) [9, 15]. Based on RCTD cell-type probability estimates, malignant cells constituted the dominant compartment in both PT and LN samples, followed by cancer-associated fibroblasts (CAFs) (Supplement Figure 1D). Spatial probability maps further revealed focally distributed immune cell populations within stromal regions (Figure 1C, Supplement Figure 1E; data shown for four the most prevalent cell types). Downstream analysis of Seurat clusters (Figure 1D, Supplement Figure 1F), revealed transcriptionally distinct clusters comprising mixed malignant and tumour microenvironment (TME) cell types.

### *TMSB4X* and *CD74* expression defines trajectory-driving clusters in tumour metastasis

Using Monocle3, we performed a trajectory analysis (Figure 2A-B, Supplement Figure 2A-F) for each sample, estimating the cellular developmental pathways by ordering cells along a pseudotemporal axis [11]. We filtered top 0.5%, 1%, 1.5%, 2% of genes with the highest Moran’s I statistic value. Across all samples, we identified eight common trajectory-driving genes (*ACTB, B2M, CD74, COL1A1, COL1A2, TMSB4X, UBC, VIM*), with *TMSB4X* and *CD74* emerging as two major trajectory drivers across all examined tissues.

**Figure 2.**
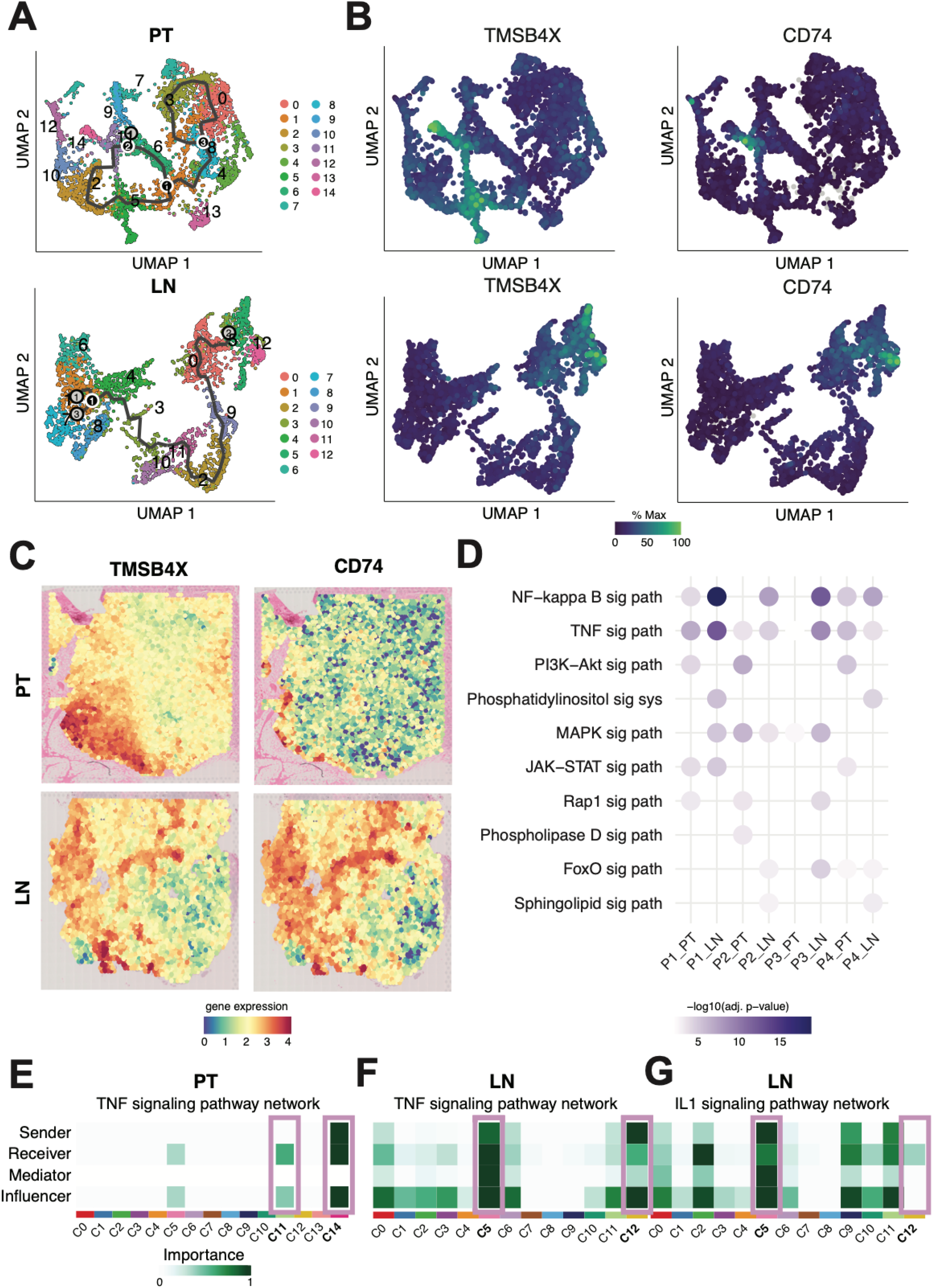
Trajectory analysis reveals NF-κB/TNF-centred signalling driven by *TMSB4X/CD74* expressing clusters. A) A Monocle 3 trajectory graph illustrating Seurat ordered clusters where nodes represent transcriptionally distinct cell states and edges depict inferred developmental transitions along differentiation lineages. B) A Monocle 3 trajectory graph illustrating genes with expression dynamics associated with cellular differentiation and developmental progression (genes with the highest Moran’s I statistic value). C) Spatial expression of genes with expression dynamics associated with cellular differentiation and developmental progression (genes with the highest Moran’s I statistic value). D) Top 5 KEGG signalling pathways within clusters expressing genes with the highest Moran’s I statistic value (*TMSB4X* and *CD74* expression). For visualization purposes multiple clusters of the same patient are merged into one. Detailed KEGG signalling is present in the Supplement Figure. E) Signalling role network graphs showing TNF and IL-1 signalling within Seurat defined clusters. Clusters with the highest expression of *TMSB4X* and *CD74* are marked. Sender - The cell type that produces and releases signalling molecules (e.g., cytokines, ligands). Receiver - The cell type that expresses the corresponding receptors and responds to the signal. Mediator - An intermediate molecule or pathway component that transmits the signal inside the receiving cell (e.g., adaptor proteins, transcription factors). Influencer - A cell or molecule that modulates the strength or direction of signalling between sender and receiver, without being the primary source or target. Figure legend: C – cluster; LN – lymph node; P – patient; PT – primary tumour; sig – signalling.

Importantly, clusters with the highest expression dynamics of *TMSB4X* and *CD74* (Patient 1 PT – c11,c14; Patient 1 LN – c5,c12; Patient 2 PT – c2; Patient 2 LN – c2,c4; Patient 3 PT – c0,c4; Patient 3 LN – c3; Patient 4 PT – c7; Patient 4 LN – c3; Figure 1D, Supplement Figure 1F) were associated with lower malignant cell type probabilities and higher probabilities of CAF and immune cell identities (Wilcoxon test p < 0.001, for all comparisons). Importantly, *TMSB4X* and *CD74* high expressing cells were spatially isolated niches in the TME localized near immune infiltrations suggesting that *TMSB4X*/*CD74*-high clusters represent transitional or communication-active states at the interface between malignant and stromal compartments (Figure 2C, Supplement Figure 2G-I).

### NF-κB/TNF-centred signalling and inflammatory reprogramming of *TMSB4X*/*CD74* clusters

Given these transcriptional and spatial features, we next asked which molecular processes underlie the behaviour of *TMSB4X*/*CD74*-high clusters. We performed a KEGG analysis of the identified DEGs [12]. The top five signalling pathways included PI3K-Akt, NF-κB, TNF, Rap1, *MAPK* signalling pathways (Supplement Figure 2J). Notably, clusters with the highest expression of *TMSB4X* and *CD74 (*Figure 1D, Supplement Figure 1E), showed predominant NF-κB and TNF signalling across all examined PTs and LN metastases (Figure 2D).

CellChat analysis examined cell-cell communication and ligand–receptor interactions within enriched signalling pathways [13]. Most trajectory driven genes were not core ligand or receptor genes in the identified KEGG pathways but were indirectly involved in related modules such as extracellular matrix (ECM), focal adhesion, ubiquitin signalling, and cytoskeleton organisation, which feed into those signalling cascades.

Cell-cell communication analysis suggested that clusters with the highest expression of *TMSB4X* and *CD74* often functioned as immune hubs (e.g., macrophages, immune cells, fibroblasts), sending, receiving, mediating, and influencing signals within the TME. These clusters were prominently engaged in NF-κB/TNF-associated signalling, as well as in ligand– receptor interaction modules annotated to PI3K-Akt and Rap1 pathways reflecting upstream growth factor- and adhesion-related ligand activity. Importantly, NF-κB/TNF signalling was rather localised to *TMSB4X*/*CD74* immune hubs in PT, but much more spread in LN mediating signalling on all clusters with additional IL-1 signalling network absent in PT (Figure 2E-G, Supplement Figure 2K-S). Conversely, MIF-CD74 interactions were not restricted to immune hubs, suggesting that *TMSB4X*, rather than *CD74*, could be a main driver in communication.

CellChat analysis of the TNF family revealed that both PTs and LN metastasis were dominated by *TNFRSF1A* signalling, consistent with canonical TNF/NF-κB activation. However, metastases, showed increased *TNFRSF1B* contribution, which is more associated with survival and immunomodulatory signalling (not shown) [16]. While TNF signalling was largely isolated in PTs, metastatic lesions showed co-activation of IL1 signalling, representing a convergent NF-κB activation [17]. This suggests metastases remodel the TNF/NF-κB axis toward a more inflammatory and pro-survival environment, potentially supporting metastatic progression and immune adaptation.

### External validation reveals myeloid to endothelial NF-κB/TNF signalling in TNBC progression

We examined 35,582 cells (Figure 3A-B, GSE180286) to evaluate *TMSB4X* and *CD74* expression patterns. Pearson correlation coefficients computed for each cell type revealed the highest correlation in myeloid cells for both PT and LN metastasis samples (Figure 3C-D).

**Figure 3.**
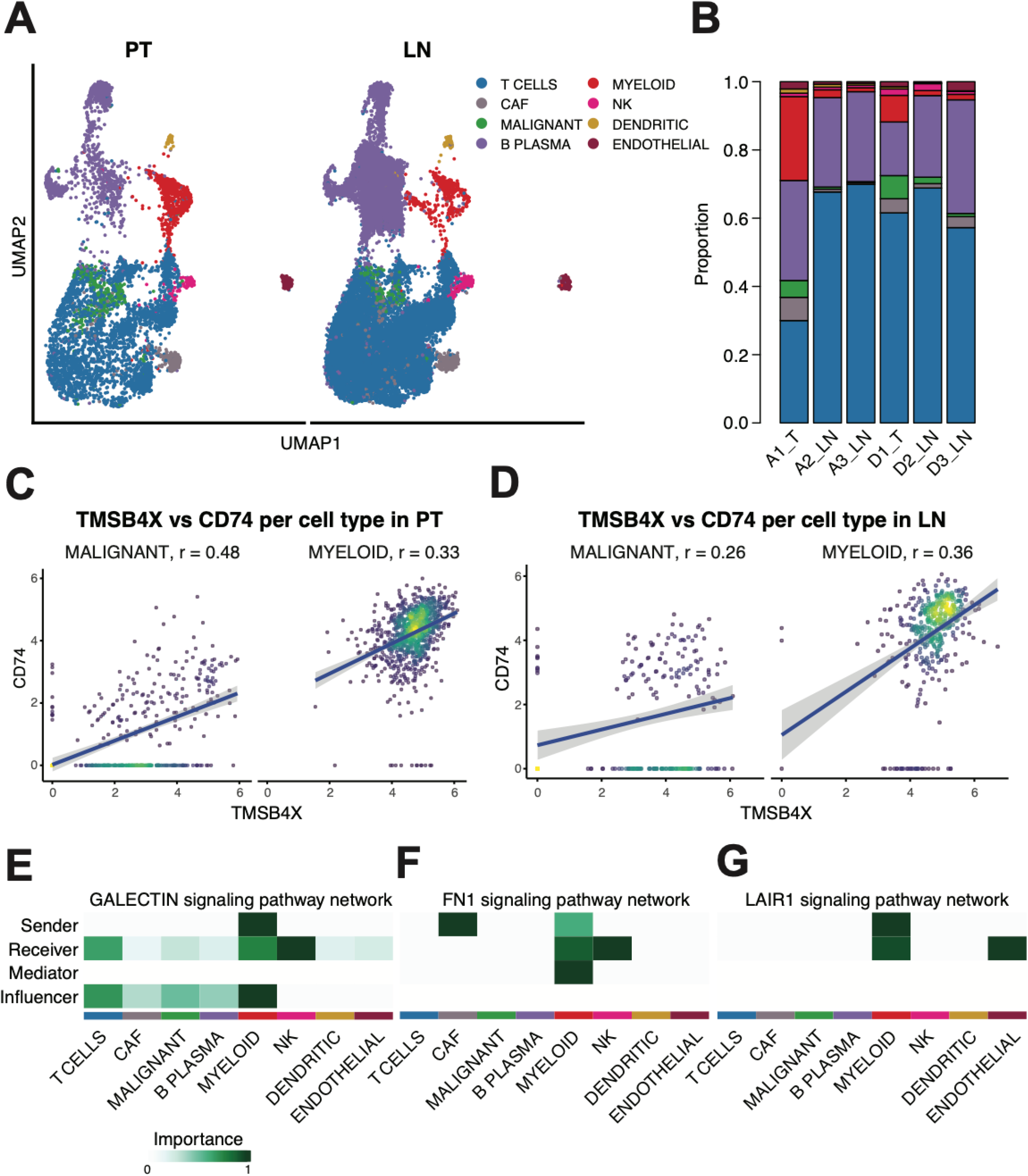
Single-cell RNA analysis reveals myeloid to endothelial NF-κB/TNF signalling in TNBC progression. A) UMAP plots of publicly available dataset of TNBC tumours and corresponding LN metastasis (GSE180286). B) Proportion of cell types of GSE180286 dataset. C-D) Gene expression correlation of TMSB4X and CD74 within different cell-types computed with Pearson correlation coefficients. Cell-types with the highest correlation scores are presented. J-K) Signalling role network graphs of major myeloid cells associated signalling pathways. Sender - The cell type that produces and releases signalling molecules (e.g., cytokines, ligands). Receiver - The cell type that expresses the corresponding receptors and responds to the signal. Mediator - An intermediate molecule or pathway component that transmits the signal inside the receiving cell (e.g., adaptor proteins, transcription factors). Influencer - A cell or molecule that modulates the strength or direction of signalling between sender and receiver, without being the primary source or target. Figure legend: CAF – cancer associated fibroblast; LN – lymph node; PT – primary tumour; T tumour

CellChat analysis in PT and LN metastases confirmed NF-κB/TNF-associated networks intersecting with *TMSB4X* and *CD74* activity. In the TME, myeloid cells acted as dominant hubs through Galectin, *CXCL, SPP1, FN1*, and *ANNEXIN* signalling, broadly influencing malignant, fibroblast, endothelial, and immune populations (Figure 3E-F, Supplement Figure A-C). Fibroblasts contributed strongly via FN1, Annexin and Laminin signalling to myeloid and NK cells (Figure 3F, Supplement Figure 3C-D). These pathways represent canonical NF-κB/TNF-driven signalling, overlapping with TMSB4X-mediated cytoskeletal remodelling and CD74-mediated antigen presentation/NF-κB activation. Counter-regulatory mechanisms such as LAIR1 signalling from myeloid to endothelial cells may temper collagen-induced NF-κB activation (Figure 3G).

In LN metastases, signalling architecture shifted, with endothelial cells emerging as major organizers. APP-driven signalling from endothelial cells targeted dendritic, myeloid, and B/plasma cells, while endothelial-endothelial communication was mediated by CD99 and SELPLG, and dendritic cells sustained CCL-mediated chemokine signalling (Supplement Figure 3E-H). This contrasted with PTs, where CAFs drove signalling through LAMIN, APP, and CD99 networks (Supplement Figure 3I-J). These findings highlight that NF-κB/TNF-driven chemokine, Galectin, and ECM pathway’s structure primary tumour communication, whereas metastatic LNs depend on endothelial hubs. TMSB4X/CD74-associated programs integrate into both settings through myeloid-endothelial crosstalk and inflammatory remodelling.

### Myeloid Cell Crosstalk with Tumour and Stromal Compartments via NF-κB–Linked and Angiogenic Signals

To validate myeloid-endothelial crosstalk in independent TNBC cohorts, we analysed scRNA BC atlas data (GSE176078, [14]) (Figure 4A). We observed the highest TMSB4X and CD74 co-expression among myeloid and epithelial (both malignant and non-malignant) cells (Figure 4B). Myeloid cells targeted endothelial compartments via CXCL, VISFATIN, VEGF, and OSM signalling, with CXCL and OSM representing canonical NF-κB/TNF-regulated pathways [18, 19] (Figure 4C-E, Supplement Figure 4A). Myeloid hubs engaged malignant, stromal, and immune cells through Galectin signalling (Figure 4F) and targeted cancer epithelial cells via EGF/DHT signalling [20], with EGF connected to downstream MAPK/PI3K-NF-κB activation [21] (Figure 4G, Supplement Figure 4B).

**Figure 4.**
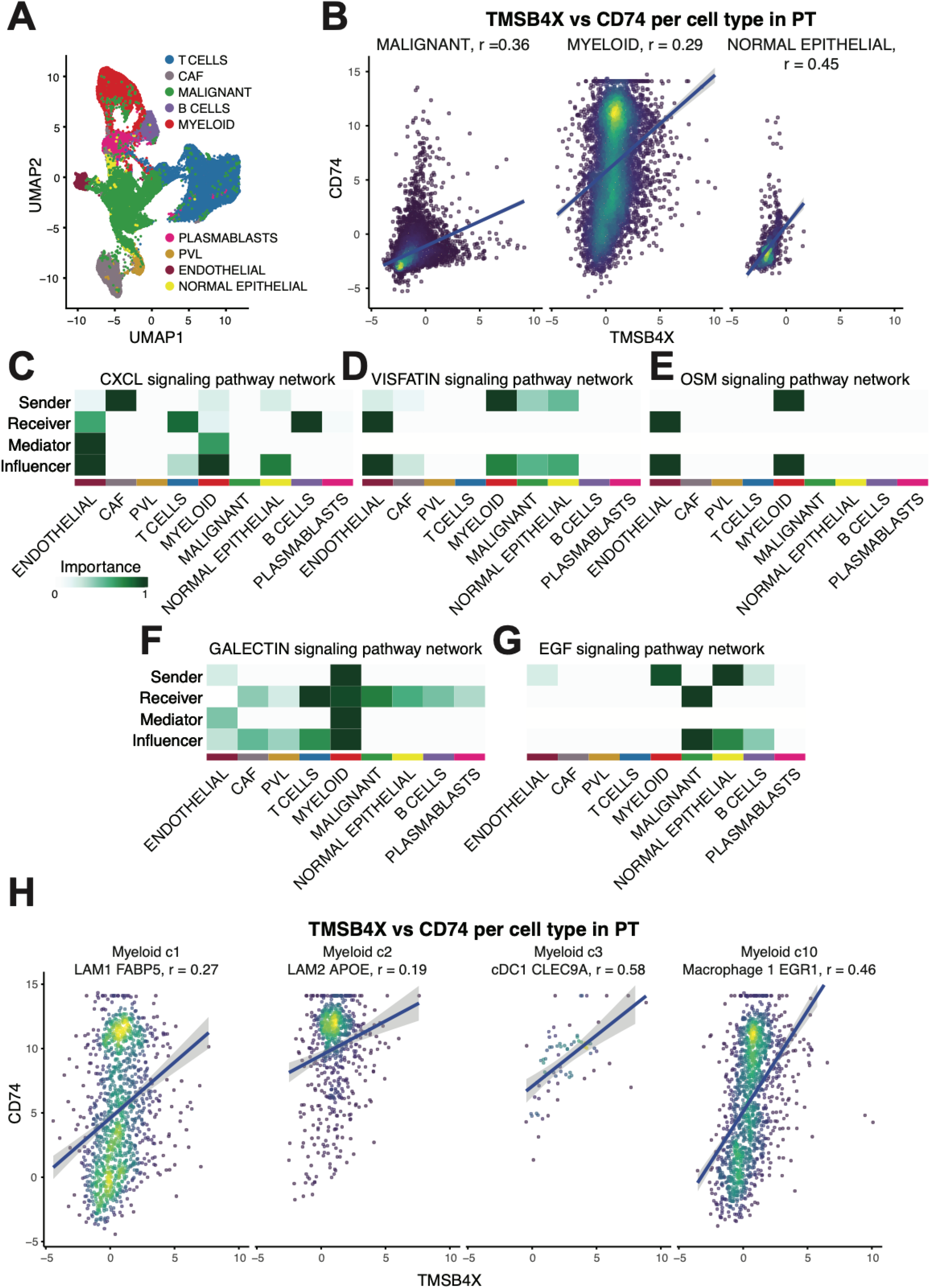
Single-Cell RNA analysis reveals myeloid cell crosstalk with tumour and stromal compartments via NF-κB–Linked and angiogenic signals. A) UMAP plot of publicly available dataset of TNBC tumours (GSE176078). B) Gene expression correlation of TMSB4X and CD74 within different cell-types computed with Pearson correlation coefficients. Cell-types with the highest correlation scores are presented. C-G) Signalling role network graphs of myeloid-endothelial cells crosstalk associated signalling pathways. Sender - The cell type that produces and releases signalling molecules (e.g., cytokines, ligands). Receiver - The cell type that expresses the corresponding receptors and responds to the signal. Mediator - An intermediate molecule or pathway component that transmits the signal inside the receiving cell (e.g., adaptor proteins, transcription factors). Influencer - A cell or molecule that modulates the strength or direction of signalling between sender and receiver, without being the primary source or target. H) Gene expression correlation of TMSB4X and CD74 for marker specific subsets of major cell-types computed with Pearson correlation coefficients Cell-types with the highest correlation scores are presented. Figure legend: CAF – cancer associated fibroblast; PVL - perivascular-like cells; PT – primary tumour.

### Endothelial-to-Myeloid and Epithelial Crosstalk Integrates Immune, Adhesion, and Metabolic Signalling Pathways

Endothelial cells influenced myeloid populations through APP, MHC-II, CD99, and Laminin signalling, with APP and MHC-II indirectly linked to NF-κB activation via antigen presentation [22, 23] (Supplement Figure 4C-F). Additional interactions with myeloid and cancer epithelial populations were mediated by Annexin, ICAM, JAM, and GAS signalling, with ICAM adhesion molecules feeding into NF-κB cascades (Supplement Figure 4G-J). In parallel, IGFBP signalling from endothelial cells affected multiple compartments, most prominently myeloid and cancer epithelial cells, and mapped preferentially to the IGF/PI3K–Akt axis rather than NF-κB/TNF (Supplementary Figure 4K-L) [24]. In contrast, normal epithelial cells broadly signalled through TNF, with the strongest effects on myeloid populations, consistent with canonical NF-κB activation (Supplementary Figure 4M). These results highlight that beyond classical NF-κB/TNF pathways (CXCL, Galectin, OSM, ICAM, Laminin, MHC-II, TNF), endocrine and metabolic signalling routes (VISFATIN, DHT, testosterone, GAS, IGFBP) diversify the regulatory network in the TNBC microenvironment.

Single-cell analysis revealed *TMSB4X*-*CD74* co-expression enrichment in specific myeloid and endothelial populations linking to distinct signalling programs (Figure 4H). LYVE1^+^ lymphatic endothelial cells, LAM1^+^/FABP5^+^ and LAM2^+^/APOE^+^ myeloid populations, EGR1+ macrophages, CLEC9A^+^ dendritic cells, and SIGLEC1^+^/CXCL10^+^ macrophages collectively engage NF-κB, PI3K-AKT, MAPK, RAS, and epithelial-mesenchymal transition (EMT) related pathways, coordinating proliferation, inflammatory responses, and metastatic remodelling.

Collectively, these results support a conceptual model of metastatic rewiring, in which localized TMSB4X-CD74 immune hubs in primary tumours give rise to expanded, multicentric and endothelial-centred inflammatory networks in lymph node metastases (Figure 5).

**Figure 5.**
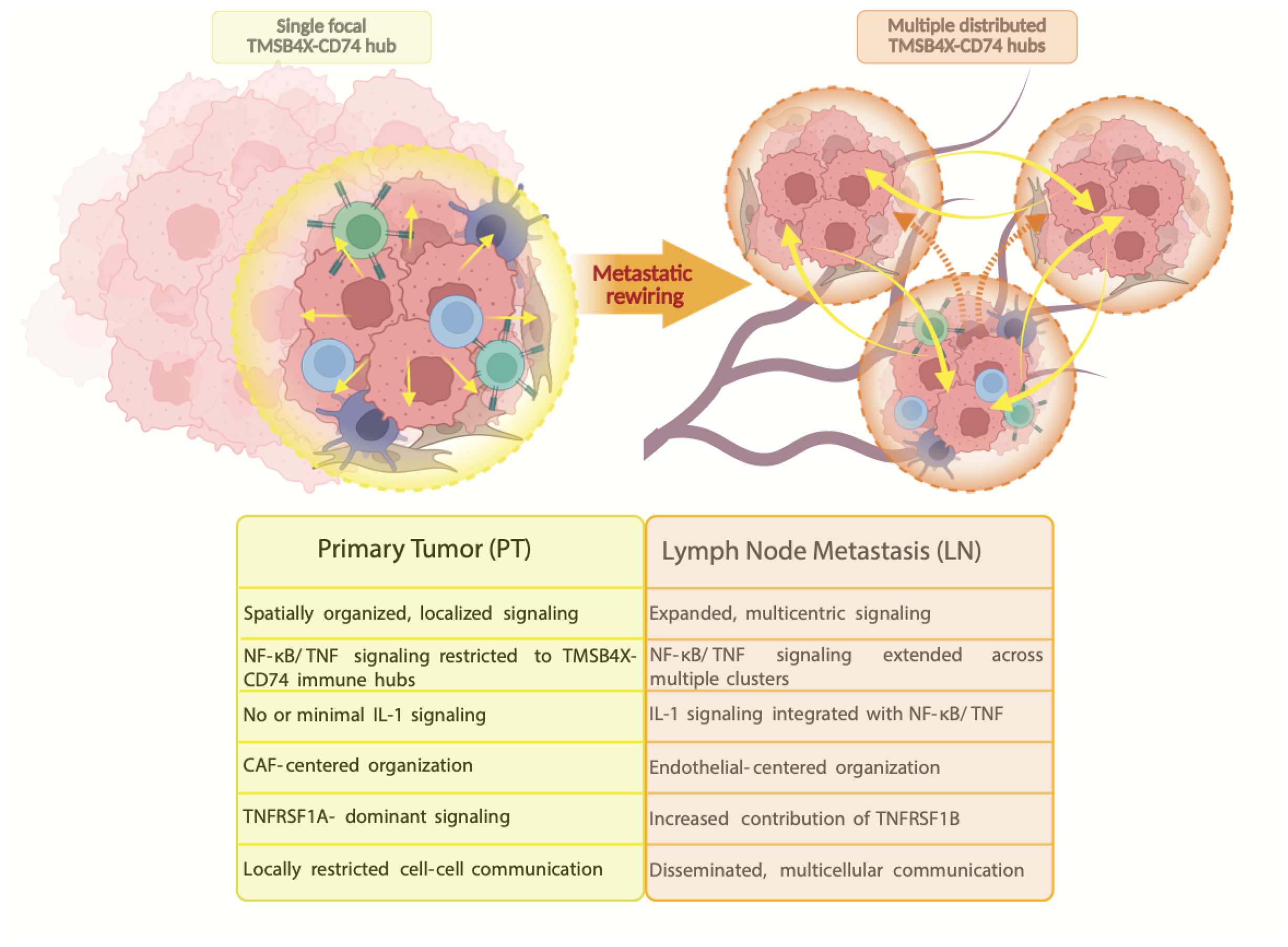
Schematic presentation of study major conclusions. Figure legend: LN – lymph node; PT – primary tumour

## DISCUSSION

In this study, we identified TMSB4X and CD74 as conserved trajectory drivers marking spatially organized immune communication hubs in TNBC. These hubs integrated actin cytoskeleton remodelling, antigen presentation, and NF-κB/TNF-centred signalling, providing a mechanistic link between cellular remodelling, immune regulation, and inflammatory communication. Validation in an independent scRNA BC atlas confirmed TMSB4X-CD74 hub architecture and revealed specific myeloid and endothelial subsets driving these interactions [14]. Our findings highlight spatially structured myeloid-endothelial crosstalk, coordinated through NF-κB/TNF signalling, that underpins TNBC metastatic ecosystems.

Our spatial transcriptomics analysis revealed that TMSB4X and CD74 are recurrently enriched in trajectory-defining clusters across patient tumours. Both genes are functionally aligned with metastasis-promoting pathways: TMSB4X regulates actin polymerization and cytoskeletal dynamics essential for migration, invasion, and EMT, while CD74 functions as the MHC-II invariant chain and MIF receptor, activating MAPK and NF-κB pathways to promote survival and inflammation [25, 26]. Critically, spatial enrichment of these genes in localized clusters indicates metastatic potential is not uniformly distributed but concentrated in defined microenvironments where immune communication and inflammatory signalling converge. This spatial organization provides additional complexity to canonical NF-κB/TNF-driven pathways in breast cancer progression [27].

A key methodological insight is the complementary value of integrating spatial and single-cell transcriptomics [28]. While Visium provided spatially resolved transcriptomes, its multicellular capture spots were dominated by fibroblasts and malignant epithelium, obscuring signals from rarer but functionally pivotal populations such as myeloid and endothelial cells. The scRNA BC atlas [14] resolved these populations with higher precision, uncovering their roles in inflammatory and angiogenic signalling, highlighting current classical Visium limitations.

Mechanistically, myeloid-endothelial crosstalk emerged as a dominant feature across primary and metastatic sites. This bidirectional network coordinates processes sustaining tumour growth and immune evasion. Endothelial cells influenced myeloid subsets through APP, MHC-II, Laminin, and ICAM signalling, while myeloid cells engaged endothelial compartments via CXCL, OSM, VEGF, and Galectin signalling. CXCL and OSM directly activate NF-κB, whereas ICAM and MHC-II mediate antigen presentation downstream of NF-κB-regulated immune responses. Concurrently, VEGF, Laminin, and APP promote angiogenesis and ECM remodelling, amplifying inflammatory signalling and establishing a vascular-immune microenvironment supporting metastatic progression. This aligns with prior work demonstrating myeloid orchestration of vascular remodelling, pre-metastatic niche formation, and immunosuppression in breast cancer [29, 30]. We extend these models by spatially mapping interactions to *TMSB4X*-*CD74* hubs where immune signalling, cytoskeletal remodelling, and vascular adaptation intersect as organizing centres of metastatic ecosystems.

Single-cell analysis pinpointed specific subsets enriched for *TMSB4X*-*CD74* co-expression that sustain tumour adaptation and spread. LYVE1^+^ lymphatic endothelial cells provide structural backbone for lymphatic dissemination, underscoring lymphatic drainage importance in early metastatic spread. Metabolically active LAM1^+^/FABP5^+^ and LAM2^+^/APOE^+^ myeloid populations couple lipid metabolism with immune regulation [14]. EGR1^+^ macrophages activate MAPK/RAS signalling and EMT induction [31], enhancing TNBC invasive properties. SIGLEC1^+^ macrophages display anti-inflammatory features [32], suggesting coexistence of opposing immune forces within the same niche. CXCL10^+^ macrophages and CLEC9A^+^ dendritic cells reinforce complexity through chemokine-mediated immune modulation and antigen presentation [33, 34]. These data indicate TMSB4X-CD74 hubs are composite niches where specialized cell types contribute synergistically to metastasis through cytoskeletal remodelling, inflammatory activation, angiogenesis, and immune evasion.

Comparison of PTs and LN metastases ST profile revealed distinct NF-κB/TNF axis rewiring. In primary tumours, signalling was restricted to localized immune hubs, whereas lymph node metastases showed expanded signalling across multiple clusters incorporating IL-1 signalling, establishing broader inflammatory networks. This IL-1 integration, a potent NF-κB activator, indicates metastatic microenvironments amplify pro-inflammatory and pro-survival pathways, facilitating immune escape, tumour outgrowth, and potentially secondary dissemination [35]. Endothelial cells assumed central organizing roles in lymph nodes compared to fibroblast dominance in primary tumours, underscoring dynamic cellular hierarchy remodelling during metastasis. These findings support the concept of metastatic niches as adaptive ecosystems where immune and vascular components are repurposed to sustain tumour persistence and progression [36].

From a translational perspective, lymph node involvement in TNBC is strongly associated with reduced overall survival and increased risk of recurrence [37, 38], underscoring the clinical importance of dissecting signalling networks that operate within metastatic niches. TMSB4X and CD74 may therefore serve as biomarkers of immune communication hubs and potential therapeutic targets, as their spatial quantification in routine FFPE biopsies could enable clinically applicable assessment of inflammatory metastatic niche activity in TNBC [39]. Targeting CD74-MIF signalling has shown promise in preclinical models [40]. Disrupting myeloid-endothelial crosstalk through CXCL/OSM/VEGF axis blockade or ICAM-mediated adhesion inhibition may attenuate NF-κB/TNF-driven inflammation and angiogenesis. Identification of specific subsets (LAM1^+^/FABP5^+^ myeloid cells, LYVE1^+^ endothelial cells) as central drivers provides rationale for therapies selectively targeting these populations. Distinct communication rewiring between primary and metastatic tumours highlights the need for stage- and site-specific therapeutic strategies. However, these hypotheses require validation in larger cohorts and experimental models, including functional perturbation studies and high-resolution spatial analyses, to confirm causal roles in metastasis.

In summary, TMSB4X-CD74-defined hubs act as spatially organized communication centres in TNBC, integrating cytoskeletal remodelling, antigen presentation, and NF-κB/TNF-driven inflammatory signalling. By coupling spatial and single-cell transcriptomics, we reveal myeloid-endothelial crosstalk as a key driver of vascular and immune adaptation during metastasis. These insights provide spatial resolution to established cancer hallmarks and suggest therapeutic opportunities targeting inflammatory, angiogenic, and cytoskeletal programs. Although targeting these pathways shows preclinical potential for overcoming immune evasion and therapy resistance, rigorous clinical evaluation and mechanistic validation are essential for translation. Future work should employ higher-resolution technologies and functional evaluation to translate these insights into effective TNBC metastasis therapies.

