## Supplemental Table and Figures for "Spatial Transcriptomics of TNBC tumours and corresponding lymph node metastasis reveals immune hubs driven by TNF/NF-κB signalling of TMSB4X and CD74 expressing cells"

| ID | Age | Smoking | Surgery | Surgery date | Adjuvant treatment | Recurrence | Survival | LN Meta | HTN | CHD | DM | GERD | AF | HT | FH |
|---|---|---|---|---|---|---|---|---|---|---|---|---|---|---|---|
| Patient 1 - FG1XW | 72 | No | MST | 09.07.2020 | Docetaxel + Cyclophosphamide (4 cycles) + RT | N | Alive | PSLN | N | Y | N | N | N | Y | N |
| Patient 2 - 21M2K | 75 | No | BCT | 12.02.2020 | Paclitaxel (9 cycles) | Y (4 months; bones) | 2022-08-20 death | SLN | Y | Y | Y | Y | N | N | N |
| Patient 3 - YX28P | 63 | No | MST | 25.11.2021 | Paclitaxel + Carboplatin (12 cycles) + RT | Y (32 months; bones) | Alive | ATN | Y | N | Y | N | N | N | Y |
| Patient 4 - 4KKBF | 73 | No | MST | 30.04.2020 | RT | N | Alive | NLF | Y | N | Y | N | Y | N | N |

Supplement Table 1 – Description of clinical and pathological data of examined patients.

ATN – axillary top node; AF – atrial fibrillation; CHD - coronary heart disease; DM – diabetes mellitus; GERD - gastroesophageal reflux disease; FH – hypercholesterolemia; HT – hypothyroidism; HTN – hypertension; MST – mastectomy; N – no; NLF – node of the lower floor; SLN – sentinel lymph node; PSLN – post sentinel lymph node; RT – radiotherapy; Y – yes;

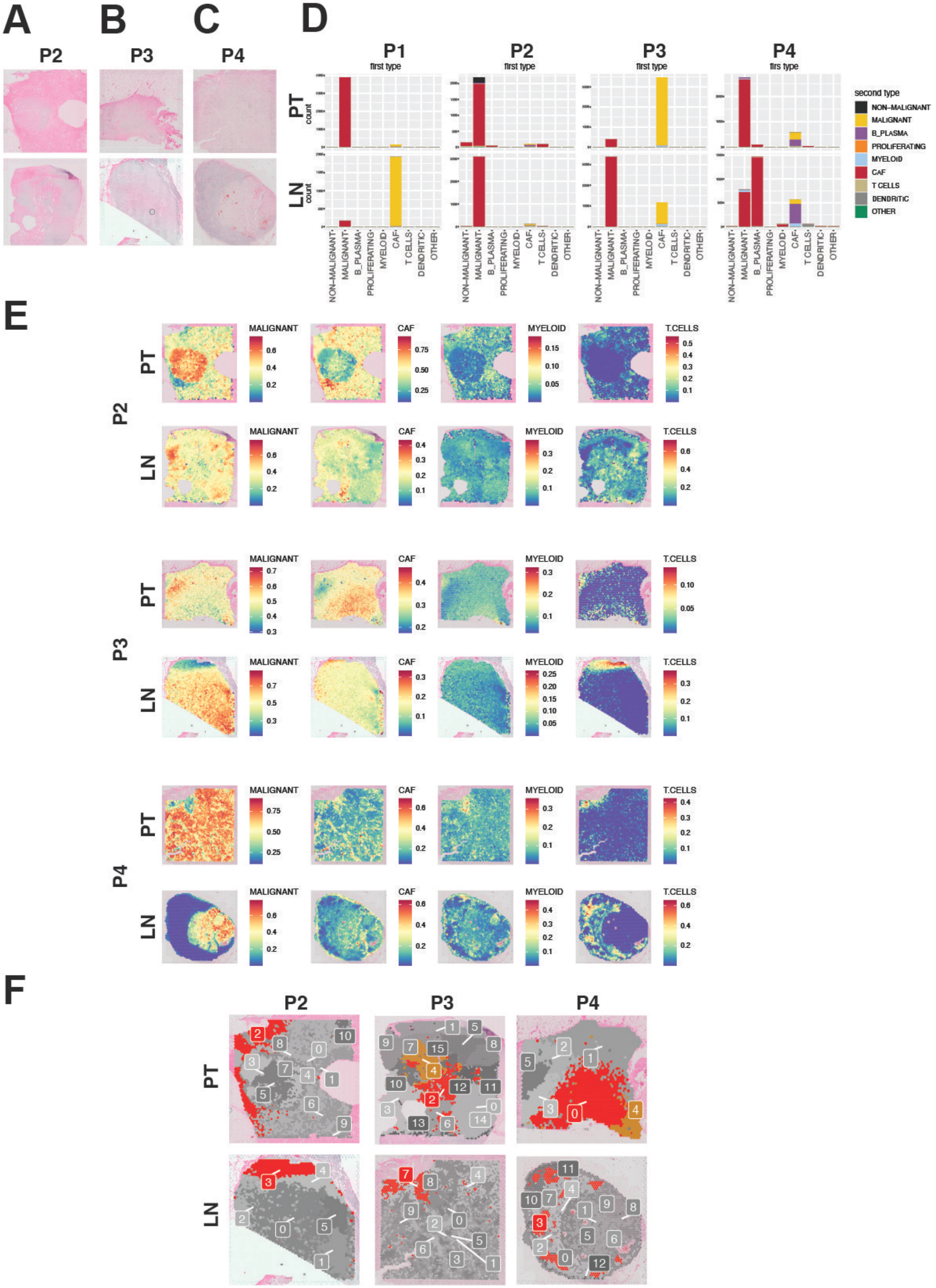


### Supplementary Figure 1

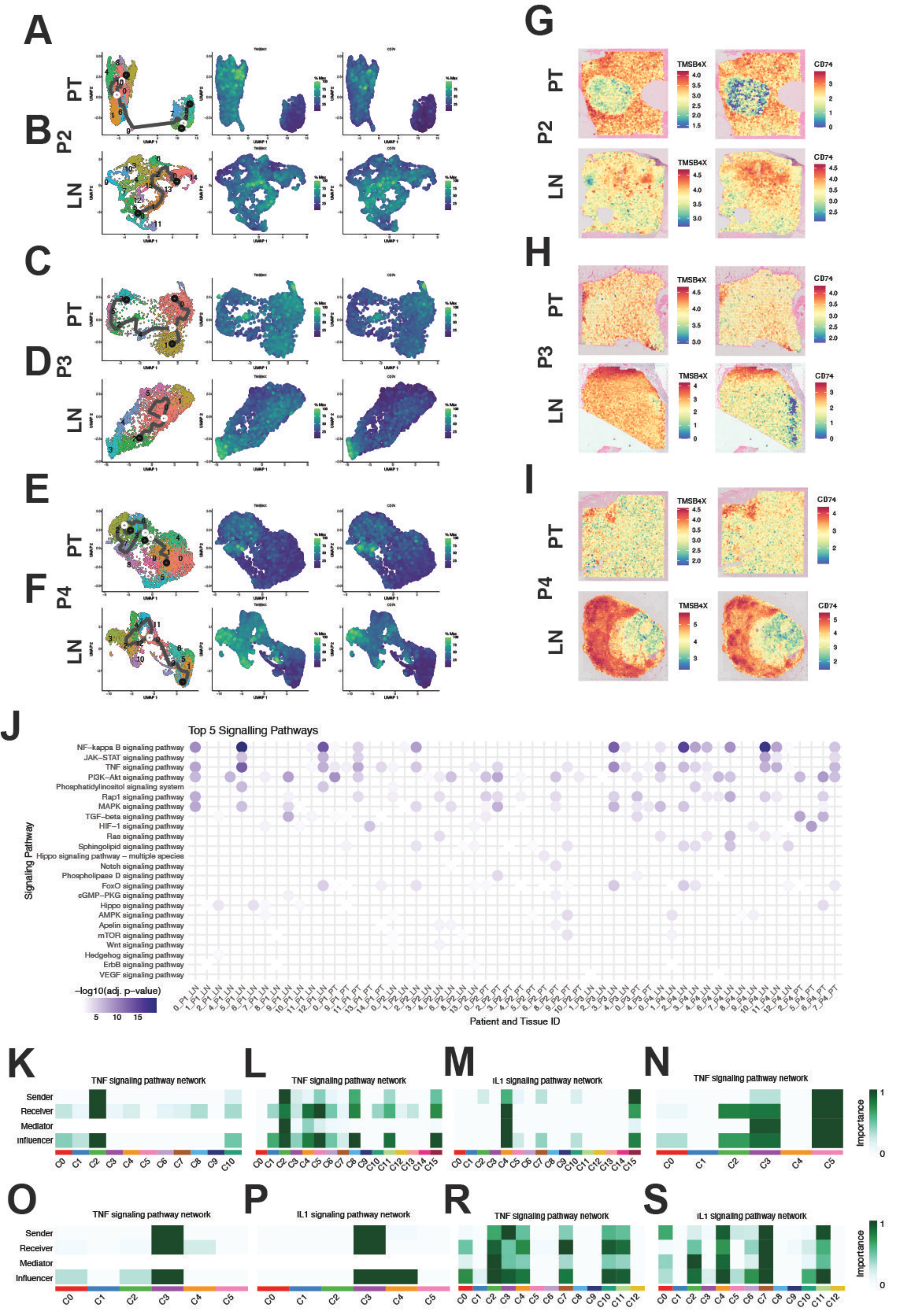


### Supplementary Figure 2

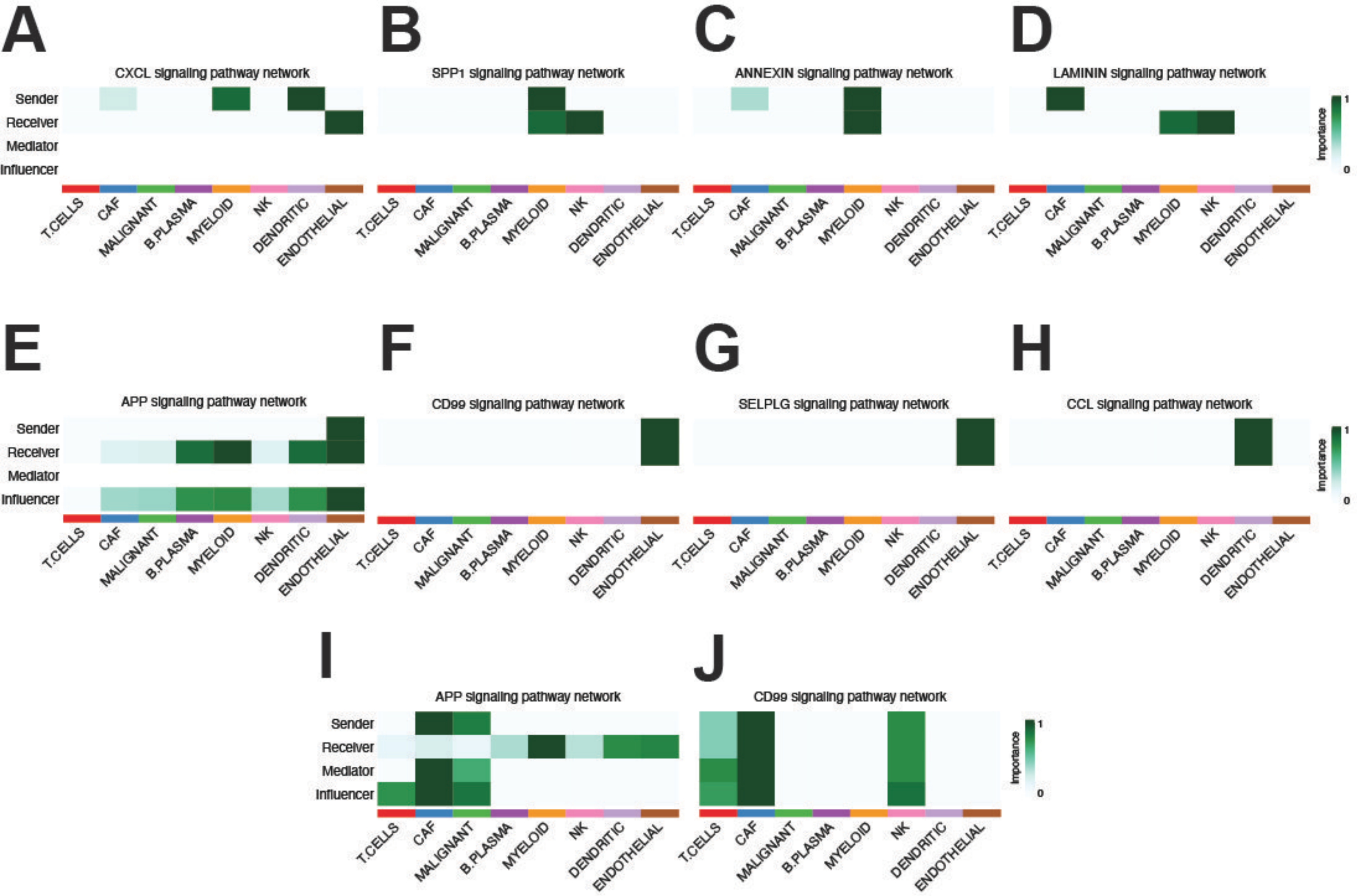


**Supplementary Figure 3**

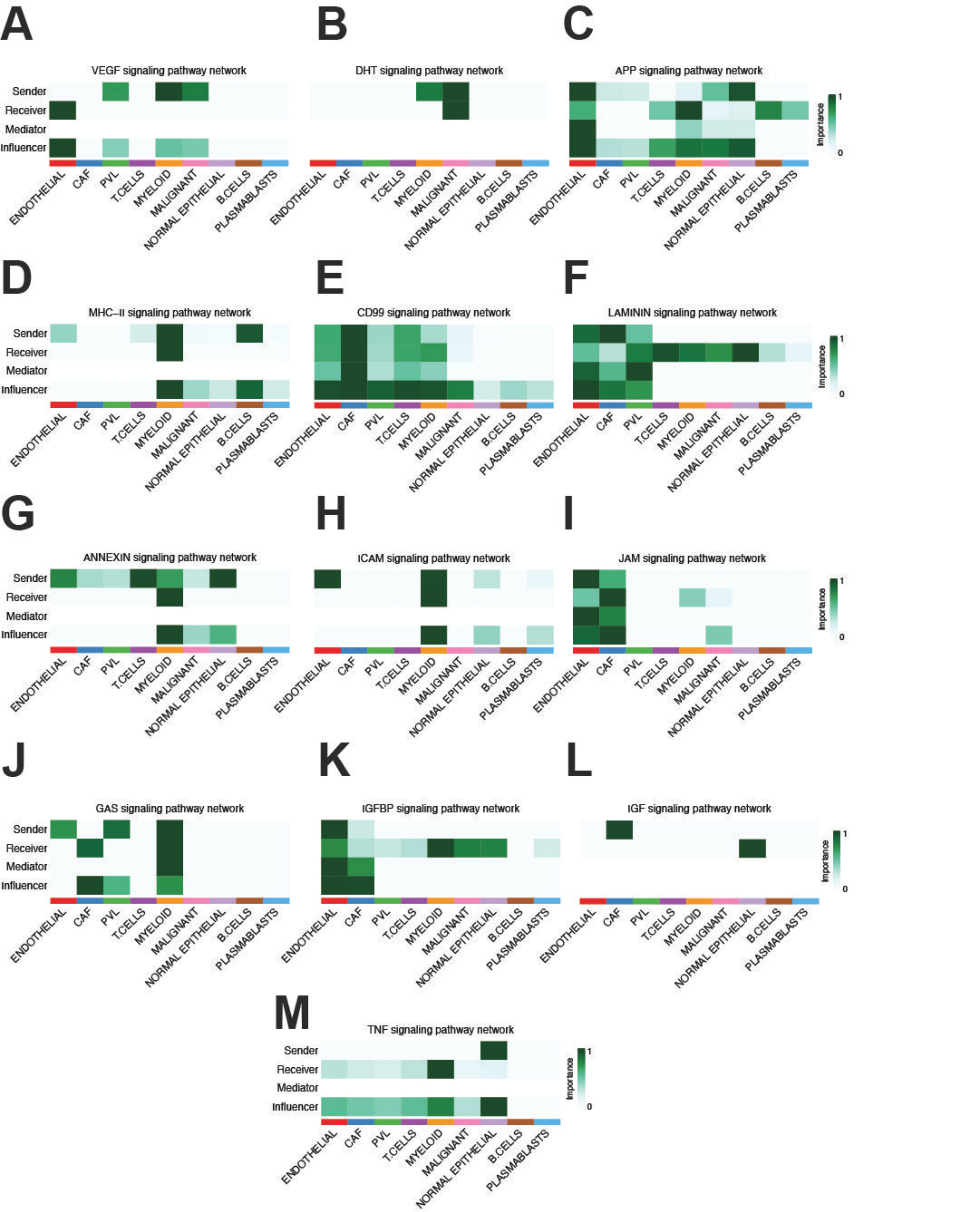


**Supplementary Figure 4**
